# Thirty two novel nsSNPs May effect on *HEXA* protein Leading to Tay-Sachs disease (TSD) Using a Computational Approach

**DOI:** 10.1101/762518

**Authors:** Tebyan A. Abdelhameed, Mohamed Mustafa Osman Fadul, Dina Nasereldin Abdelrahman Mohamed, Amal Mohamed Mudawi, Sayaf Kamal Khalifa Fadul Allah, Ola Ahmed Elnour Ahmed, Sogoud Mohammednour Idrees Mohammeddeen, Aya Abdelwahab Taha khairi, Soada Ahmed Osman, Ebrahim Mohammed Al-Hajj, Mustafa Elhag, Mohamed Ahmed Hassan Salih

## Abstract

**Background:** Genetic polymorphisms in the *HEXA* gene are associated with a neurodegenerative disorder called Tay-Sachs disease (TSD) (GM2 gangliosidosis type 1). This study aimed to predict the possible pathogenic SNPs of this gene and their impact on the protein using different bioinformatics tools.

**Methods:** SNPs retrieved from the NCBI database were analyzed using several bioinformatics tools. The different algorithms collectively predicted the effect of single nucleotide substitution on both structure and function of the hexosaminidase A protein.

**Results:** Fifty nine mutations were found to be highly damaging to the structure and function of the *HEXA* gene protein.

**Conclusion:** According to this study, thirty two novel nsSNP in *HEXA* are predicted to have possible role in Tay-Saches Disease using different bioinformatics tools. Our findings could help in genetic study and diagnosis of Tay-Saches Disease.

## INTRODUCTION

Tay-Sachs disease (TSD -[OMIM* 272800] also called GM2 gangliosidosis type 1 is an autosomal recessive lysosomal storage disorder which mainly affects Ashkenazi Jewish (AJ) population with low occurrences in non-AJ populations [1–5]. It’s incidence is one in 100,000 live births (carrier frequency of about one in 250) [6]. Deficiency of ß-hexosaminidase-A (Hex-A) isozyme which cleaves the terminal N-acetyl hexosamine residues of GM2 ganglioside is the cause of the TSD. This deficiency leads to accumulation of GM2 ganglioside which is caused by mutations in *HEXA* gene the highest concentration of GM2 ganglioside found in neuronal cells and they are the main glycolipids of neuronal cell’s plasma membranes responsible of the normal cellular activities. Patients with HexA deficiency suffer from GM2 ganglioside accumulates inside neural’s lysosomes. Therefore, HexA deficiency primarily affects the nervous system [1, 2, 4, 6–14].

According to the disease onset age, TSD is differentiated into three types which are: acute infantile, juvenile, and late-onset. Acute infantile TSD is the most common type. It causes progressive weakness and motor skills lost in the infected infants. This progression occurs between the ages of six months to three years. In addition, the infected infant progressively suffers from diminished attentiveness and an exaggerated startle response. As TSD continues to destroy the brain infant suffers from seizures, blindness, and death which usually occurs before four years of age [8, 9, 11, 13–15].

The human *HEXA* gene [OMIM *606869] encodes the alpha subunit of the lysosomal hexosaminidase A isozyme (HexA) which is located on chromosome 15 q23-q24 and contains 14 exons. More than two hundred mutations have been reported in the *HEXA* gene to cause TSD and its variants. These mutations include single base substitutions, small deletion, duplications, insertions splicing alterations, complex gene rearrangement, partial large duplications, partial deletion, splicing mutations, nonsense mutations and missense mutations that lead to the disruption of transcription, translation, folding, dimerization of monomers and catalytic dysfunction of HexA protein [4, 8, 11, 15, 16].

Many studies through different population demonstrate that the molecular base of *HEXA* gene still is unidentified [3, 8, 17–20]. Therefore, this study aims to recognize the possible pathogenic SNPs in *HEXA* gene using bioinformatics prediction tools to define its role of the SNPs on the structure, function, and regulation of their corresponding protein.

Recently, bioinformatics tools enabled drawing out very valuable results from large amounts of raw data like analysis of gene and protein expression, comparison of genetic data (evolution), modeling of 3D structure from sequence of gene or protein based on homologous. It also assists in prediction of damaging SNPs and its relationship with diseases [21]. Our study aimed to define the impact of *HEXA* SNPs on the structure and function of *HEXA* protein using bioinformatics softwares which may have a significant role in disease susceptibility. The usage of the bioinformatics approach has a potent impact on the identification of provider SNPs since it is easy and less costly, less time consuming and can facilitate future genetic studies. This study regarded as the first study that determine the possible damaging *HEXA* gene SNPs using bioinformatics tools which could be useful for further genetic studies.

## MATERIALS AND METHODS

Data on *HEXA* gene was obtained from the national center for biological information (NCBI) website (https://www.ncbi.nlm.nih.gov/) and the SNPs (single nucleotide polymorphisms) information was retrieved from NCBI SNPs database dbSNP (https://www.ncbi.nlm.nih.gov/snp/). The gene ID and sequence was obtained from Uiprot (https://www.uniprot.org/). Analysis of the SNPs was done according to figure 1.

**Figure (1):**
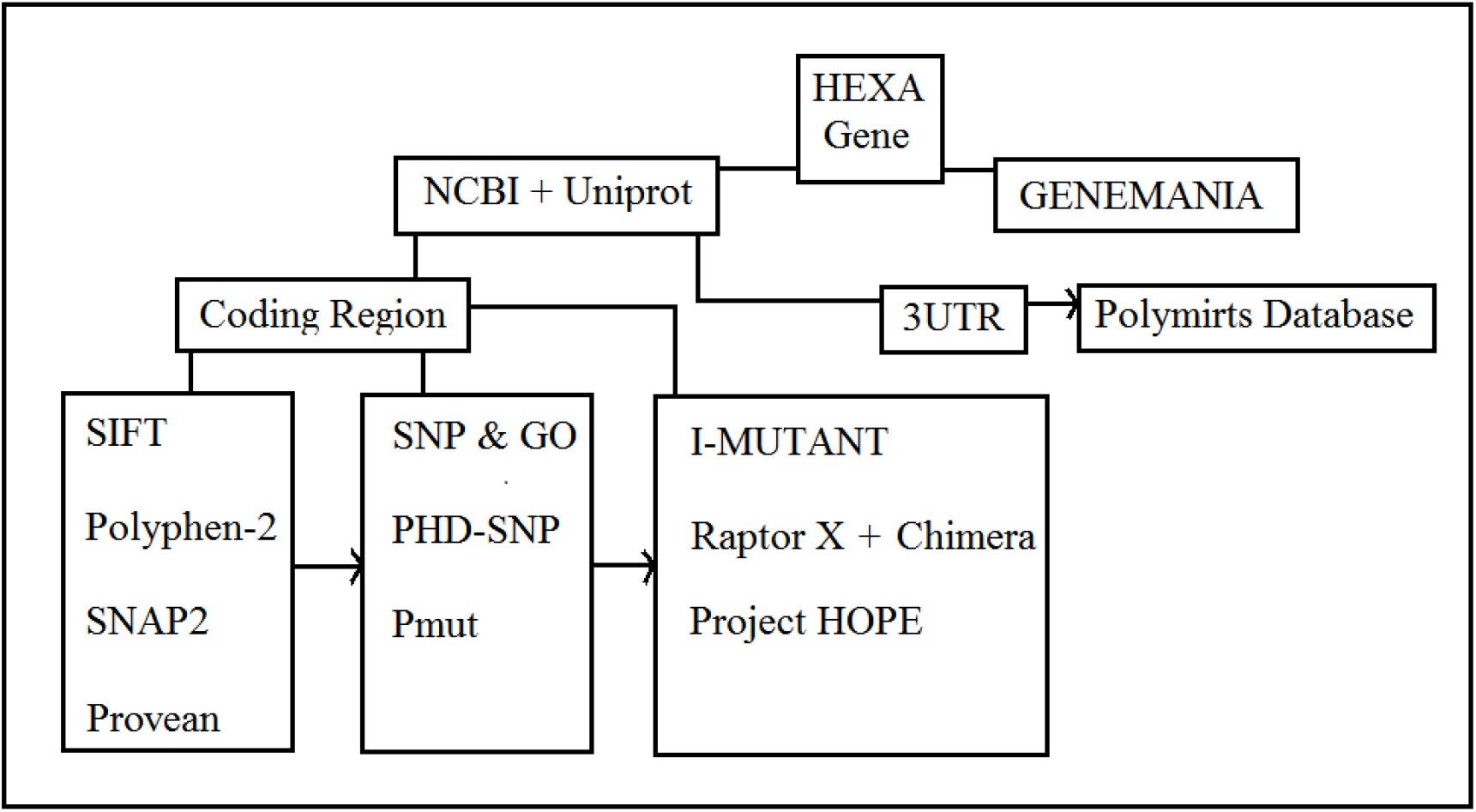
Diagram demonstrates *HEXA* gene work flow using different bioinformatics tools

### SIFT

(Sorting Intolerant from Tolerant) is a sequence homology-based tool that sorts intolerant from tolerant amino acid substitutions and predicts whether an amino acid substitution in a protein will have a phenotypic outcome, considering the position at which the mutation occurred and the type of amino acid change. When a protein sequence submitted, SIFT chooses related proteins and obtains an alignment of these proteins with the query. Based on the change on the type of amino acids appearing at each position in the alignment, SIFT calculates the probability that amino acid at a position is tolerated conditional on the most frequent amino acid being tolerated. If this normalized value is less than a cutoff, the substitution is predicted to be deleterious. SIFT scores <0.05 are predicted by the algorithm to be intolerant or deleterious amino acid substitutions, whereas scores >0.05 are considered tolerant [22]. (http://sift.bii.a-star.edu.sg/)

### PolyPhen-2

Polymorphism Phenotyping v.2 available at (http://genetics.bwh.harvard.edu/pph2/) is a tool that predicts the possible effect of an amino acid substitution on the structure and function of a human protein by using simple physical and comparative considerations. The submission of the sequence allows querying for a single individual amino acid substitution or a coding, non-synonymous SNP annotated in the SNP database. It calculates position-specific independent count (PSIC) scores for each of the two variants and computes the difference of the PSIC scores of the two variants. The higher a PSIC score difference, the higher the functional effect of a particular amino acid substitution is likely to have. PolyPhen scores were designated as probably damaging (0.95–1), possibly damaging (0.7–0.95), and benign (0.00– 0.31) [23].

### Provean

Protein Variation Effect Analyzer software available at (http://provean.jcvi.org/index.php) it predicts whether an amino acid substitution has an impact on the biological function of the protein. Provean is useful in filtrating sequence variants to identify non-synonymous variants that are predicted to be functionally important [24].

### SNAP2

Available at (https://rostlab.org/owiki/index.php/Snap2), SNAP2 is a trained classifier that is based on a machine learning device called “neural network”. It distinguishes between effect and neutral variants/non-synonymous SNPs by taking a variety of sequence and variant features in considerations. The most important input signal for the prediction is the evolutionary information taken from an automatically generated multiple sequence alignment. Also, structural features such as predicted secondary structure and solvent accessibility are considered. If available also annotation (i.e. known functional residues, pattern, regions) of the sequence or close homologs are pulled in. SNAP2 has persistent two-state accuracy (effect/neutral) of 82 % [25].

### PHD-SNP

Prediction of human Deleterious Single Nucleotide Polymorphisms is available at (http://snps.biofold.org/phd-snp/phd-snp.html). The working principle is Support Vector Machines (SVMs) based method trained to predict disease-associated nsSNPs using sequence information. The related mutation is predicted as disease-related (Disease) or as neutral polymorphism (Neutral) [26].

### SNP and GO

It is a server for the prediction of single point protein mutations likely to be involved in the causing of diseases in humans.[26] (https://snps-and-go.biocomp.unibo.it/snps-and-go/)

### Pmut

Available at (http://mmb.irbbarcelona.org/PMut/),Pmut is based on the use of different kinds of sequence information to label mutations, and neural networks to process this information. It provides a very simple output: a yes/no answer and a reliability index [27].

### IMUTANT

Imutant is a suit of support vector machine based predictors which integrated into a unique web server. It offers the opportunity to predict automatically protein stability changes upon single site mutations starting from protein sequence alone or protein structure if available. Also, it gives the possibility to predict human deleterious SNPs from the protein sequence alone. It is vailable at (http://gpcr.biocomp.unibo.it/~emidio/I-Mutant3.0/old/IntroI-Mutant3.0_help.html) [28].

### Project Hope

Online software is available at: (http://www.cmbi.ru.nl/hope/method/). It is a web service where the user can submit a sequence and mutation. The software collects structural information from a series of sources, including calculations on the 3D protein structure, sequence annotations in UniProt and prediction from other software. It combines this information to give analysis for the effect of a certain mutation on the protein structure. HOPE will show the effect of that mutation in such a way that even those without a bioinformatics background can understand it. It allows the user to submit a protein sequence (can be FASTA or not) or an accession code of the protein of interest. In the next step, the user can indicate the mutated residue with a simple mouse click. In the final step, the user can simply click on one of the other 19 amino acid types that will become the mutant residue, and then full report well is generated [29].

### UCSF Chimera (University of California at San Francisco)

UCSF Chimera (https://www.cgl.ucsf.edu/chimera/) is a highly extensible program for interactive visualization and analysis of molecular structures and related data, including density maps, supramolecular assemblies, sequence alignments, docking results, trajectories, and conformational ensembles. High-quality images and animations can be generated. Chimera includes complete documentation and several tutorials. Chimera is developed by the Resource for Biocomputing, Visualization, and Informatics (RBVI), supported by the National Institutes of Health (P41-GM103311).[30]

### PolymiRTS

PolymiRTS is a software used to predict 3UTR (un-translated region) polymorphism in microRNAs and their target sites available at (http://compbio.uthsc.edu/miRSNP/). It is a database of naturally occurring DNA variations in mocriRNAs (miRNA) seed region and miRNA target sites. MicroRNAs pair to the transcript of protein coding genes and cause translational repression or mRNA destabilization. SNPs in microRNA and their target sites may affect miRNA-mRNA interaction, causing an effect on miRNA-mediated gene repression, PolymiRTS database was created by scanning 3UTRs of mRNAs in human and mouse for SNPs in miRNA target sites. Then, the effect of polymorphism on gene expression and phenotypes are identified and then linked in the database. The PolymiRTS data base also includes polymorphism in target sites that have been supported by a variety of experimental methods and polymorphism in miRNA seed regions. [31]

### GeneMANIA

It is gene interaction software that finds other genes which is related to a set of input genes using a very large set of functional association data. Association data include protein and genetic interactions, pathways, co-expression, co-localization and protein domain similarity. GeneMANIA also used to find new members of a pathway or complex, find additional genes you may have missed in your screen or find new genes with a specific function, such as protein kinases. available at (https://genemania.org/) [32].

## RESULTS

### The functional effect of Deleterious and damaging nsSNPs of *HEXA* by SIFT, PolyPhen-2, Provean, and SNAP2

**Table (1):** Prediction of functional effect of Deleterious and damaging nsSNPs by SIFT, Polyphen-2, PROVEAN and SNAP2

| Mutation | SIFT<br>PRE | SIFT<br>SCO | POLYPHEN PRE | POLYPHEN<br>SCO | PROVEAN<br>SCO | PROVEAN<br>PRE | SNAP2<br>SCO | SNAP2<br>PRE |
| --- | --- | --- | --- | --- | --- | --- | --- | --- |
| G511D | D | 0 | PROB DAMAGE | 1 | -6.394 | Deleterious | 76 | effect |
| G511V | D | 0 | PROB DAMAGE | 1 | -8.221 | Deleterious | 61 | effect |
| R499H | D | 0 | PROB DAMAGE | 1 | -4.642 | Deleterious | 97 | effect |
| R499C | D | 0 | PROB DAMAGE | 1 | -7.427 | Deleterious | 97 | effect |
| A496P | D | 0 | PROB DAMAGE | 1 | -4.242 | Deleterious | 11 | effect |
| W485R | D | 0 | PROB DAMAGE | 1 | -12.998 | Deleterious | 97 | effect |
| E482K | D | 0 | PROB DAMAGE | 1 | -3.714 | Deleterious | 98 | effect |
| P475R | D | 0 | PROB DAMAGE | 1 | -8.356 | Deleterious | 76 | effect |
| W474C | D | 0 | PROB DAMAGE | 1 | -12.209 | Deleterious | 83 | effect |
| R472S | D | 0 | PROB DAMAGE | 1 | -5.648 | Deleterious | 71 | effect |
| E462V | D | 0 | PROB DAMAGE | 1 | -6.621 | Deleterious | 85 | effect |
| G454A | D | 0 | PROB DAMAGE | 1 | -5.675 | Deleterious | 79 | effect |
| G454S | D | 0 | PROB DAMAGE | 1 | -5.675 | Deleterious | 98 | effect |
| G454R | D | 0 | PROB DAMAGE | 1 | -7.566 | Deleterious | 93 | effect |
| P439L | D | 0 | PROB DAMAGE | 1 | -9.569 | Deleterious | 50 | effect |
| W420C | D | 0 | PROB DAMAGE | 1 | -12.44 | Deleterious | 94 | effect |
| S417Y | D | 0 | PROB DAMAGE | 1 | -4.879 | Deleterious | 76 | effect |
| D378Y | D | 0 | PROB DAMAGE | 1 | -8.787 | Deleterious | 9 | effect |
| I335N | D | 0 | PROB DAMAGE | 1 | -6.341 | Deleterious | 62 | effect |
| C328Y | D | 0 | PROB DAMAGE | 1 | -10.723 | Deleterious | 49 | effect |
| D322N | D | 0 | PROB DAMAGE | 1 | -4.877 | Deleterious | 78 | effect |
| D322Y | D | 0 | PROB DAMAGE | 1 | -8.731 | Deleterious | 88 | effect |
| G321E | D | 0 | PROB DAMAGE | 1 | -7.803 | Deleterious | 86 | effect |
| G321R | D | 0 | PROB DAMAGE | 1 | -7.803 | Deleterious | 89 | effect |
| L319H | D | 0 | PROB DAMAGE | 1 | -6.661 | Deleterious | 57 | effect |
| H318L | D | 0 | PROB DAMAGE | 1 | -10.742 | Deleterious | 88 | effect |
| Y316C | D | 0 | PROB DAMAGE | 1 | -7.45 | Deleterious | 48 | effect |
| Y298C | D | 0 | PROB DAMAGE | 1 | -8.772 | Deleterious | 52 | effect |
| D258H | D | 0 | PROB DAMAGE | 1 | -6.827 | Deleterious | 96 | effect |
| R252H | D | 0 | PROB DAMAGE | 1 | -4.877 | Deleterious | 53 | effect |
| R252C | D | 0 | PROB DAMAGE | 1 | -7.803 | Deleterious | 68 | effect |
| G250V | D | 0 | PROB DAMAGE | 1 | -8.778 | Deleterious | 67 | effect |
| G250S | D | 0 | PROB DAMAGE | 1 | -5.852 | Deleterious | 41 | effect |
| R247W | D | 0 | PROB DAMAGE | 1 | -7.803 | Deleterious | 89 | effect |
| A246T | D | 0 | PROB DAMAGE | 1 | -3.768 | Deleterious | 31 | effect |
| A246S | D | 0 | PROB DAMAGE | 1 | -2.726 | Deleterious | 13 | effect |
| F211S | D | 0 | PROB DAMAGE | 1 | -7.73 | Deleterious | 88 | effect |
| S210F | D | 0 | PROB DAMAGE | 1 | -5.823 | Deleterious | 80 | effect |
| H204R | D | 0 | PROB DAMAGE | 1 | -7.797 | Deleterious | 89 | effect |
| W203G | D | 0 | PROB DAMAGE | 1 | -12.67 | Deleterious | 77 | effect |
| H202R | D | 0 | PROB DAMAGE | 1 | -7.797 | Deleterious | 86 | effect |
| N199S | D | 0 | PROB DAMAGE | 1 | -4.873 | Deleterious | 27 | effect |
| N199I | D | 0 | PROB DAMAGE | 1 | -8.772 | Deleterious | 71 | effect |
| K197T | D | 0 | PROB DAMAGE | 1 | -5.848 | Deleterious | 95 | effect |
| Y180H | D | 0 | PROB DAMAGE | 1 | -4.448 | Deleterious | 95 | effect |
| H179L | D | 0 | PROB DAMAGE | 1 | -10.6 | Deleterious | 76 | effect |
| R178H | D | 0 | PROB DAMAGE | 1 | -4.815 | Deleterious | 94 | effect |
| R178C | D | 0 | PROB DAMAGE | 1 | -7.708 | Deleterious | 96 | effect |
| D175A | D | 0 | PROB DAMAGE | 1 | -7.708 | Deleterious | 88 | effect |
| R170Q | D | 0 | PROB DAMAGE | 1 | -3.854 | Deleterious | 96 | effect |
| R170W | D | 0 | PROB DAMAGE | 1 | -7.708 | Deleterious | 98 | effect |
| R166H | D | 0 | PROB DAMAGE | 1 | -4.818 | Deleterious | 18 | effect |
| R166C | D | 0 | PROB DAMAGE | 1 | -7.708 | Deleterious | 37 | effect |
| I161T | D | 0 | PROB DAMAGE | 1 | -4.534 | Deleterious | 43 | effect |
| R137Q | D | 0 | PROB DAMAGE | 1 | -3.654 | Deleterious | 72 | effect |
| S129F | D | 0 | PROB DAMAGE | 1 | -3.307 | Deleterious | 48 | effect |
| L127R | D | 0 | PROB DAMAGE | 1 | -5.481 | Deleterious | 95 | effect |
| Y116F | D | 0 | PROB DAMAGE | 1 | -3.682 | Deleterious | 60 | effect |
| Y68D | D | 0 | PROB DAMAGE | 1 | -8.377 | Deleterious | 65 | effect |

### Functional analysis of *HEXA* gene using diseased related softwares (PhD-SNP, SNPs & GO and PMut)

**Table (2):**
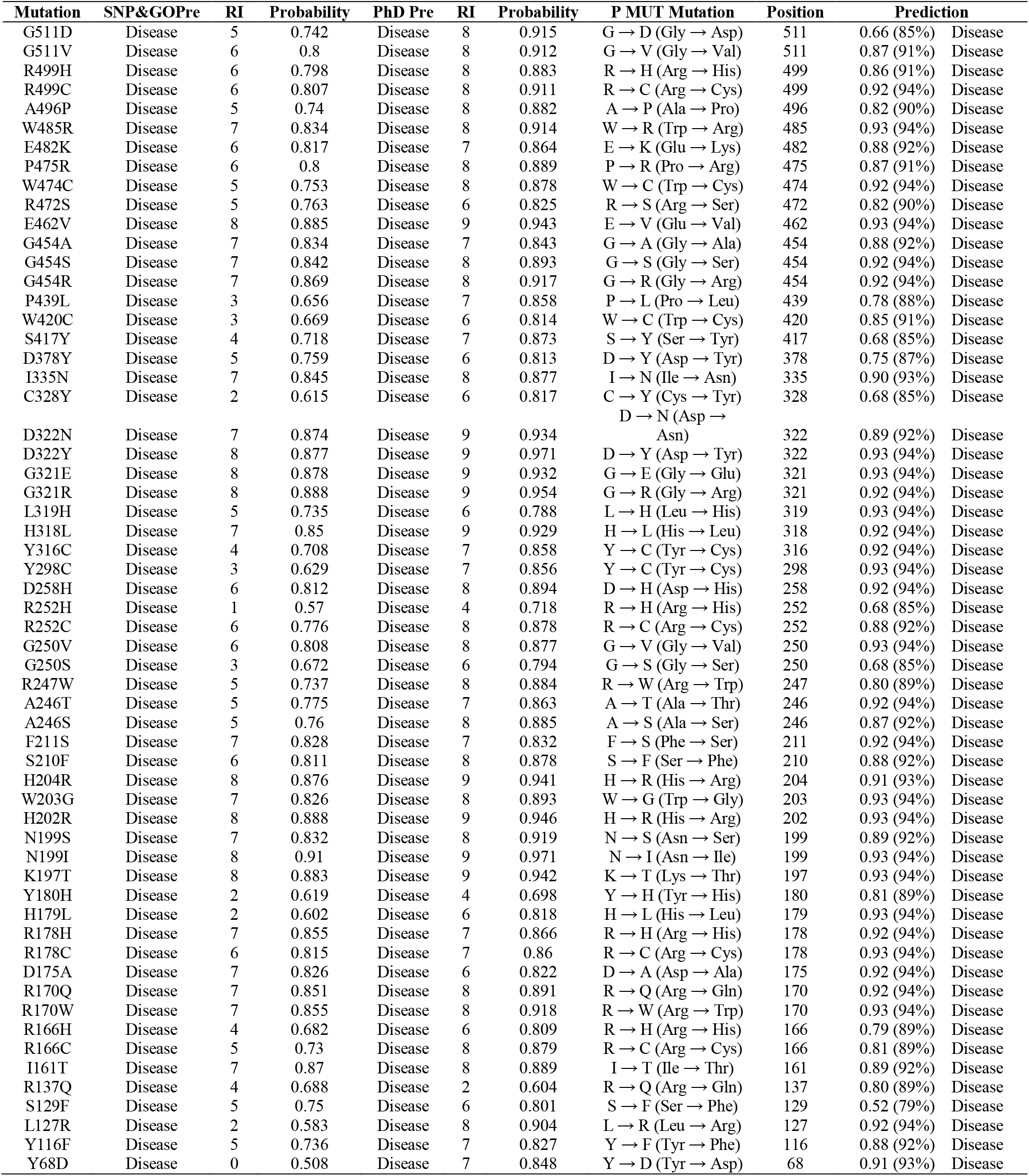
Prediction of Disease Related and pathological effect of nsSNPs by PhD-SNP, SNPs & GO and PMut:

### Prediction of Change in *HEXA* Stability due to Mutation Using I-Mutant 3.0 Server

**Table (3):** Prediction of nsSNPs Impact on Protein structure Stability by I-Mutant

Prediction of Change in *HEXA* Stability due to Mutation Using I-Mutant 3.0 Server **Table (3):** Prediction of nsSNPs Impact on Protein structure Stability by I-Mutant
| dbSNPs# | Amino Acid Change | I-Mutant PRE | I Mutant SCO |
| --- | --- | --- | --- |
| rs1448744870 | G511D | Decrease | -0.77 |
| rs1448744870 | G511V | Decrease | -0.35 |
| rs121907956 | R499H | Decrease | -1.27 |
| rs121907966 | R499C | Decrease | -0.82 |
| rs891448877 | A496P | Increase | -0.24 |
| rs121907968 | W485R | Decrease | -1.22 |
| rs121907952 | E482K | Decrease | -0.89 |
| rs758829206 | P475R | Decrease | -1.02 |
| rs121907981 | W474C | Decrease | -1.21 |
| rs369117208 | R472S | Decrease | -1.36 |
| rs863225434 | E462V | Increase | 0.2 |
| rs1229811721 | G454A | Decrease | -0.26 |
| rs121907978 | G454S | Decrease | -0.97 |
| rs121907978 | G454R | Decrease | -0.14 |
| rs1438534723 | P439L | Decrease | -0.11 |
| rs121907958 | W420C | Decrease | -1.72 |
| rs1321597225 | S417Y | Increase | -0.05 |
| rs772807186 | D378Y | Increase | -0.03 |
| rs1431312050 | I335N | Decrease | -1.68 |
| rs761861041 | C328Y | Increase | -0.23 |
| rs772180415 | D322N | Decrease | -0.53 |
| rs772180415 | D322Y | Increase | 0.03 |
| rs1316178162 | G321E | Decrease | -0.53 |
| rs556877212 | G321R | Decrease | -0.41 |
| rs1265331930 | L319H | Decrease | -2.32 |
| rs766622566 | H318L | Increase | 0.38 |
| rs1222599742 | Y316C | Decrease | -1.15 |
| rs1396944040 | Y298C | Decrease | -1.1 |
| rs121907971 | D258H | Decrease | -0.55 |
| rs762255098 | R252H | Decrease | -1.42 |
| rs566580738 | R252C | Decrease | -1.07 |
| rs121907959 | G250V | Decrease | -0.55 |
| rs1057521137 | G250S | Decrease | -1.3 |
| rs121907970 | R247W | Decrease | -0.49 |
| rs758166013 | A246T | Decrease | -0.69 |
| rs758166013 | A246S | Decrease | -0.87 |
| rs121907974 | F211S | Decrease | -1.64 |
| rs121907961 | S210F | Increase | 0.15 |
| rs121907976 | H204R | Decrease | -0.22 |
| rs1002712424 | W203G | Decrease | -2.34 |
| rs761536872 | H202R | Increase | -0.14 |
| rs1345279330 | N199S | Decrease | -0.55 |
| rs1345279330 | N199I | Increase | 0.77 |
| rs121907973 | K197T | Decrease | -0.38 |
| rs28941771 | Y180H | Decrease | -1.1 |
| rs747372270 | H179L | Increase | -0.01 |
| rs28941770 | R178H | Decrease | -1.1 |
| rs121907953 | R178C | Decrease | -0.86 |
| rs1057519460 | D175A | Decrease | -0.82 |
| rs121907957 | R170Q | Decrease | -1.07 |
| rs121907972 | R170W | Decrease | -0.44 |
| rs398123442 | R166H | Decrease | -1.43 |
| rs759487463 | R166C | Decrease | -1.08 |
| rs765136263 | I161T | Decrease | -2.07 |
| rs769724246 | R137Q | Decrease | -0.64 |
| rs1295567201 | S129F | Increase | 0.32 |
| rs121907975 | L127R | Decrease | -1.46 |
| rs1282031681 | Y116F | Decrease | -0.37 |
| rs760202948 | Y68D | Decrease | -1.11 |

### Modeling of amino acid substitution effects on *HEXA* protein structure using Chimera and Project Hope

**Figure 2:**
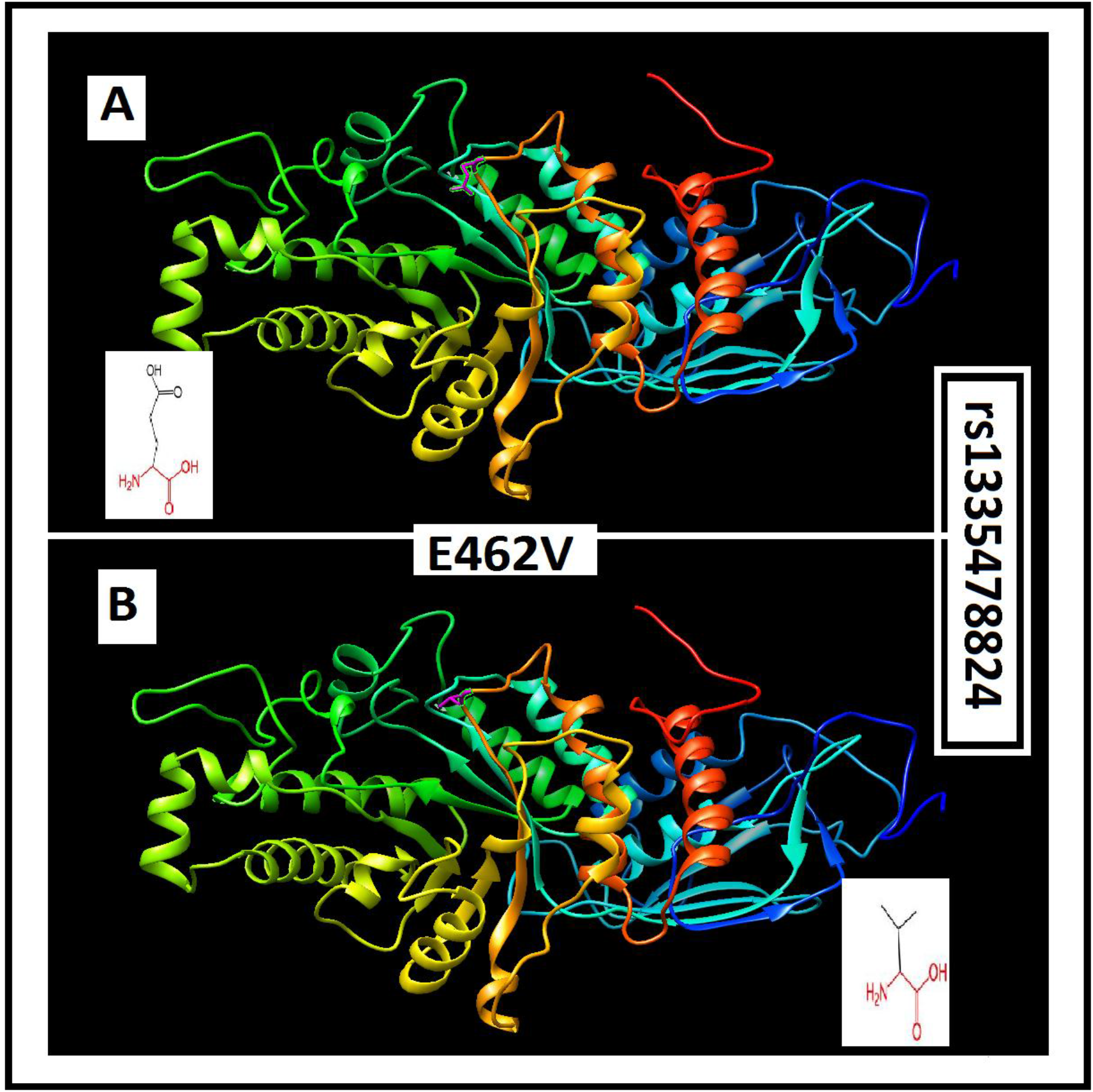
rs1335478824 (E462V): Effect of change in the amino acid position 462 from to Glutamic Acid to Valine on the *HEXA* protein 3D structure using Project Hope and Chimera Softwares.

**Figure 3:**
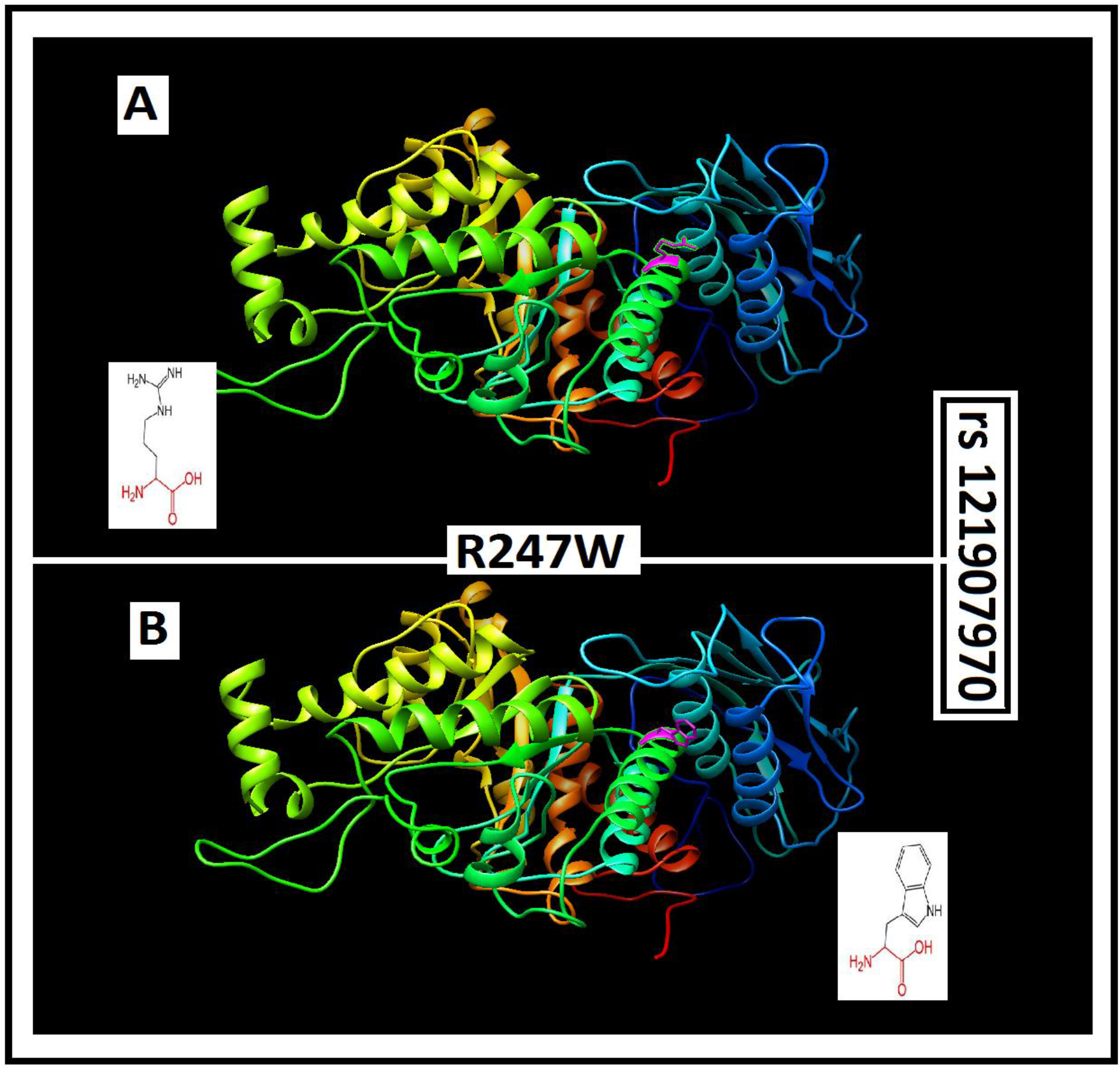
rs121907970 (R247W): Effect of change in the amino acid position 247 from Arginine to Tryptophan on the *HEXA* protein 3D structure using Project Hope and Chimera Softwares.

**Figure 4:**
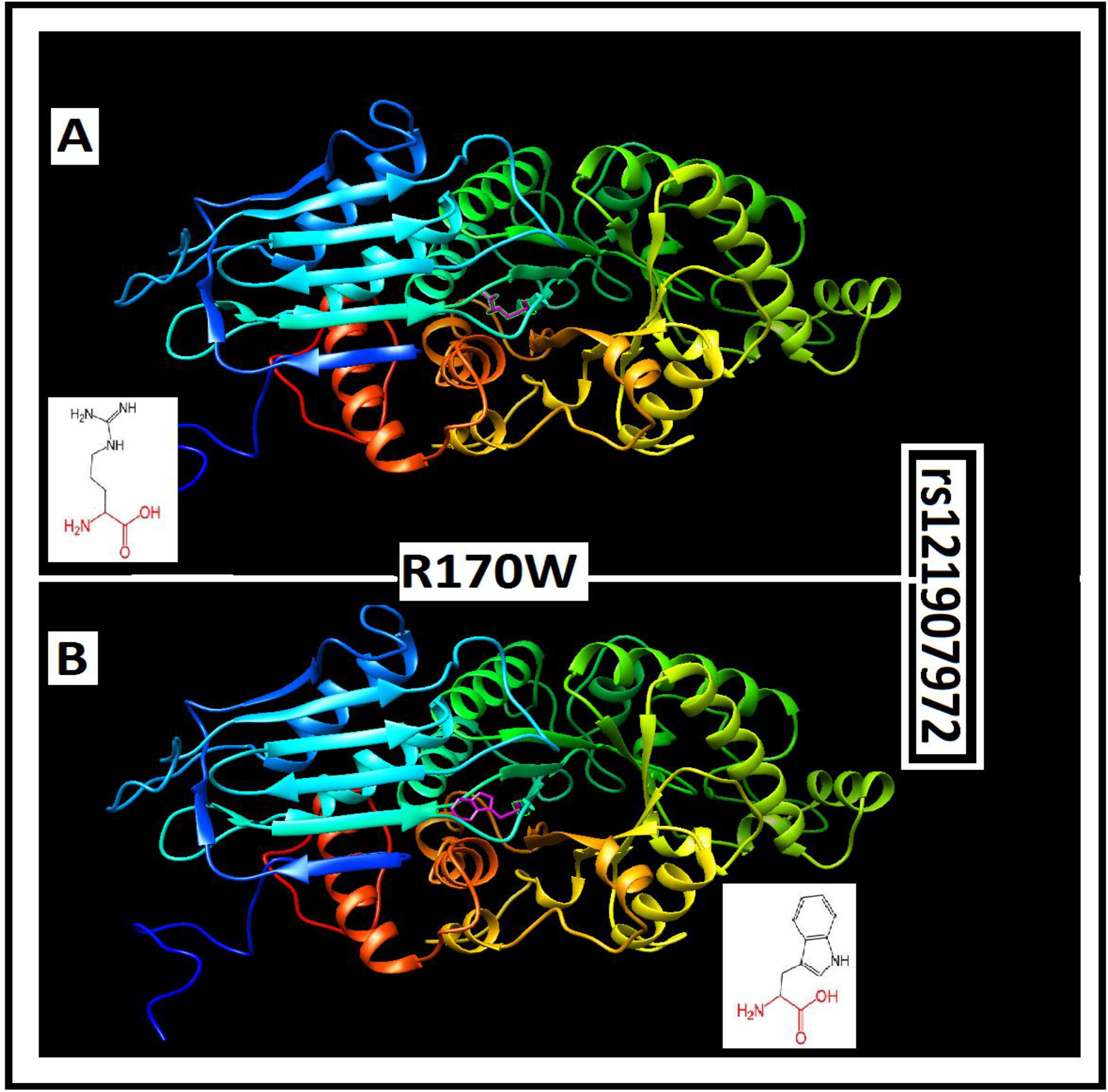
rs121907972 (R170W): Effect of change in the amino acid position 170 from Arginine toTryptophan on the *HEXA* protein 3D structure using Project Hope and Chimera Software.

**Figure 4:**
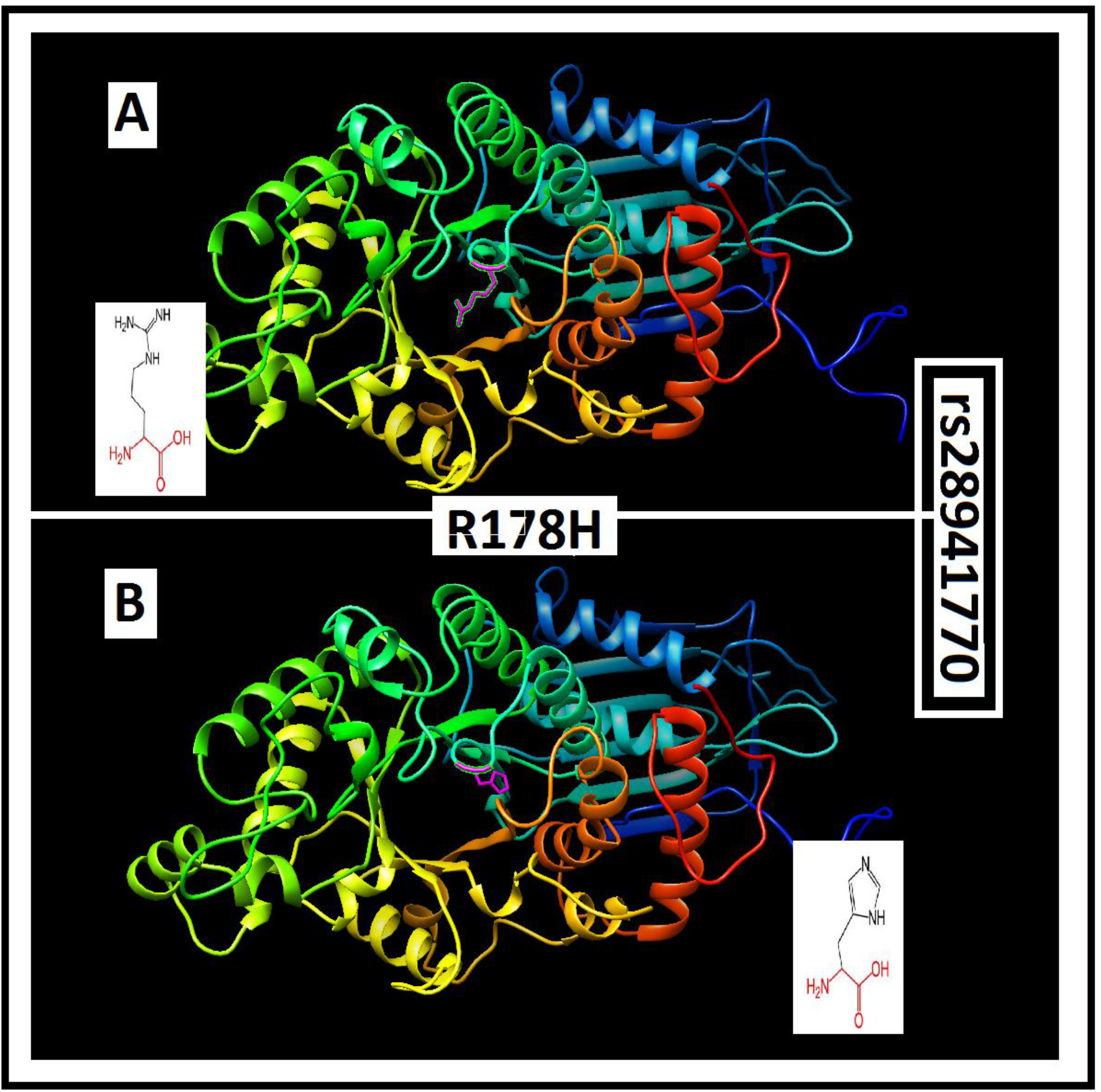
rs28941770 (R178W): Effect of change in the amino acid position 178 from Arginine to Tryptophan on the *HEXA* protein 3D structure using Project Hope and Chimera Software.

### Interactions of *HEXA* gene with other Functional Genes illustrated by GeneMANIA

**Figure 5:**
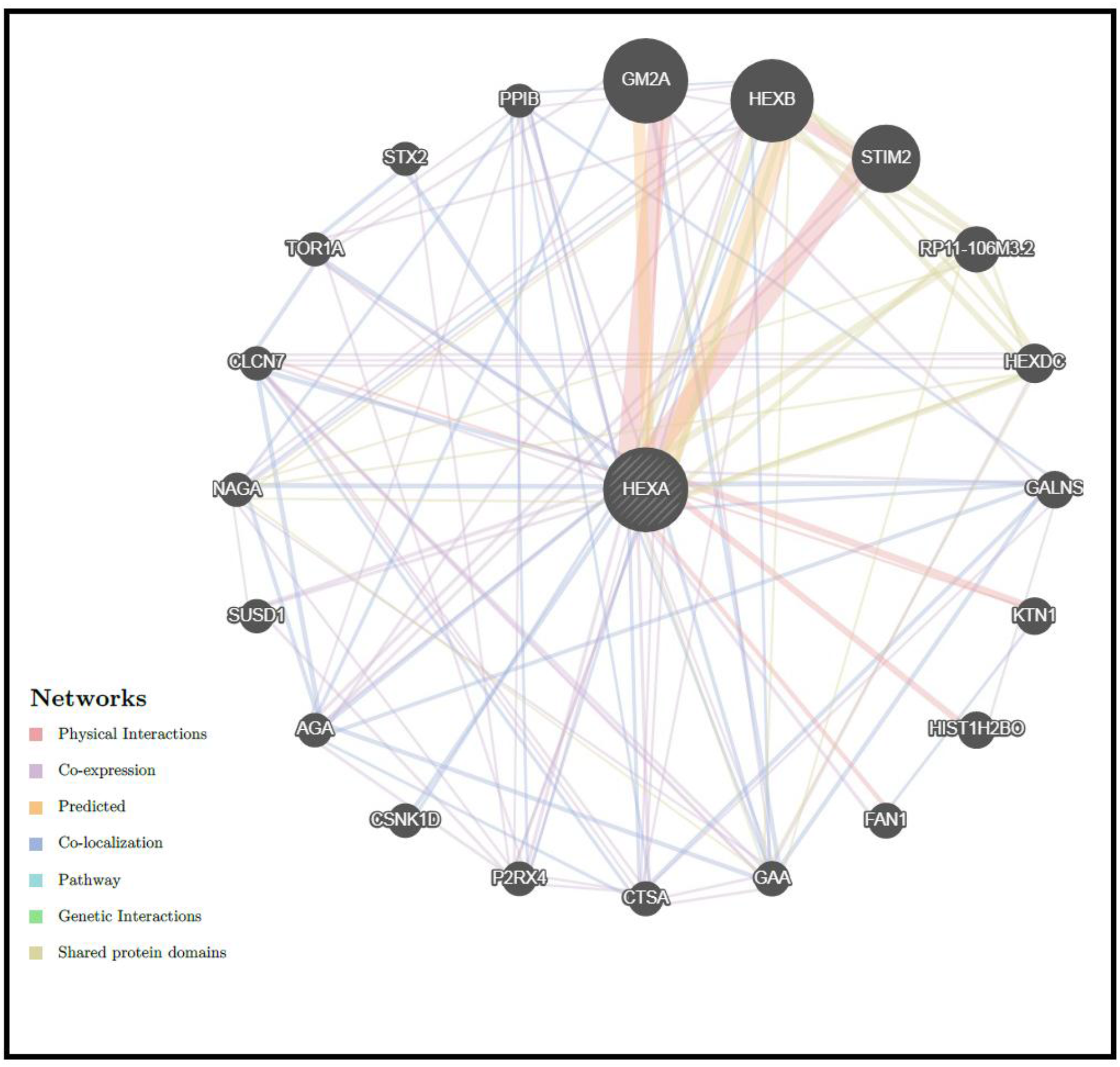
*HEXA* Gene Interactions and network predicted by GeneMania.

**Table (4):**
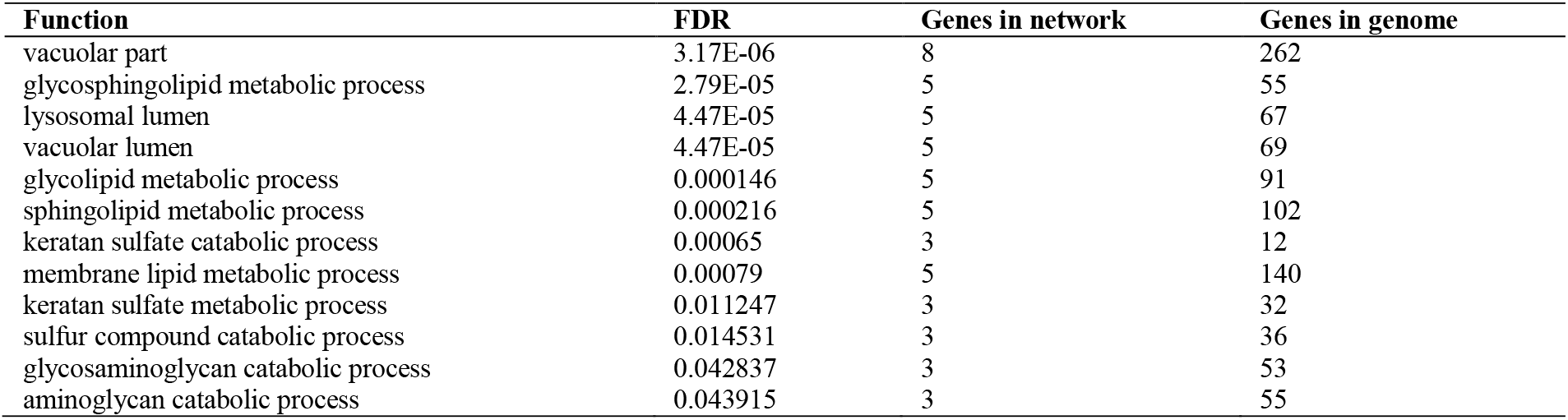
The *HEXA* gene functions and its appearance in network and genome

**Table (5):**
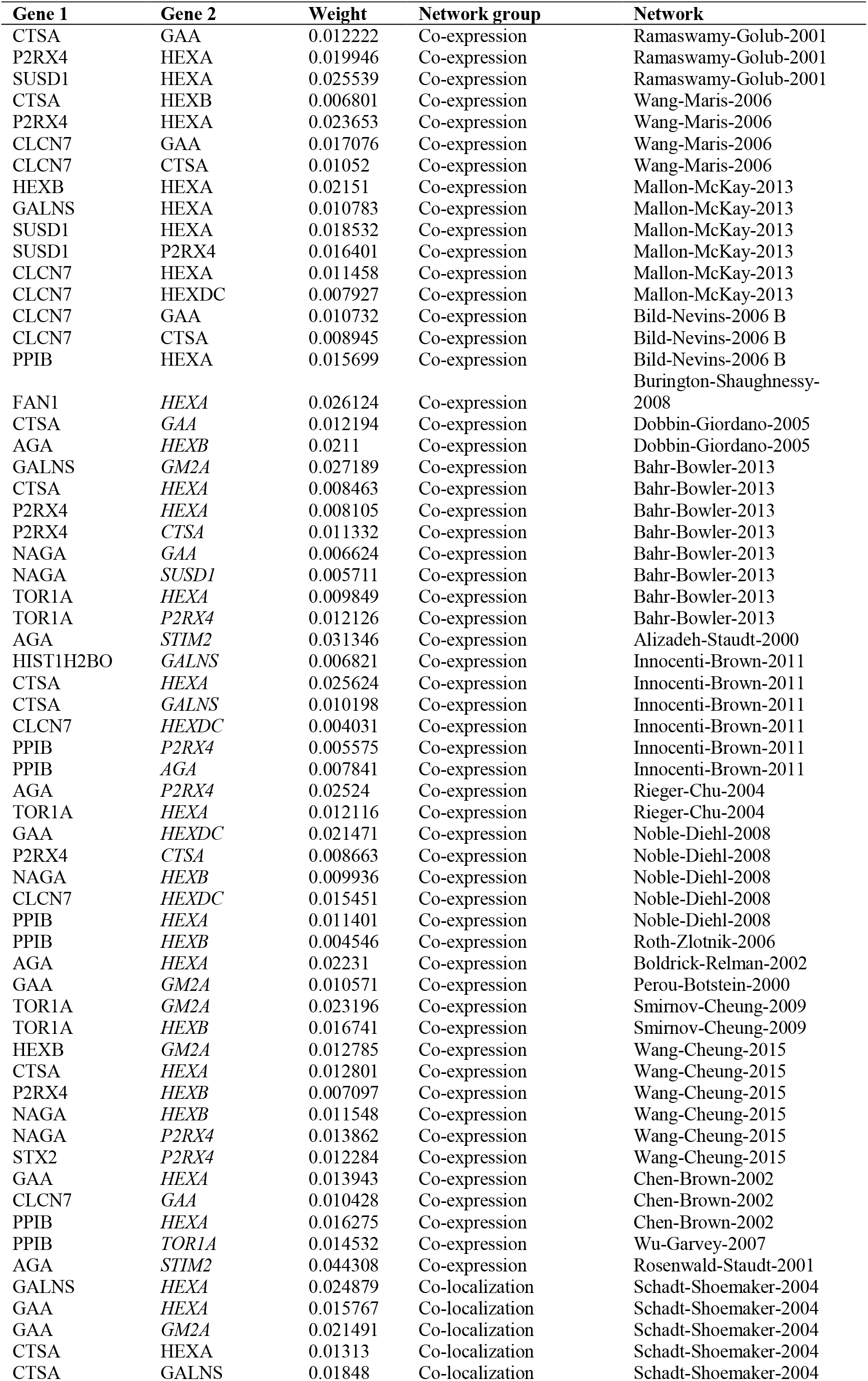
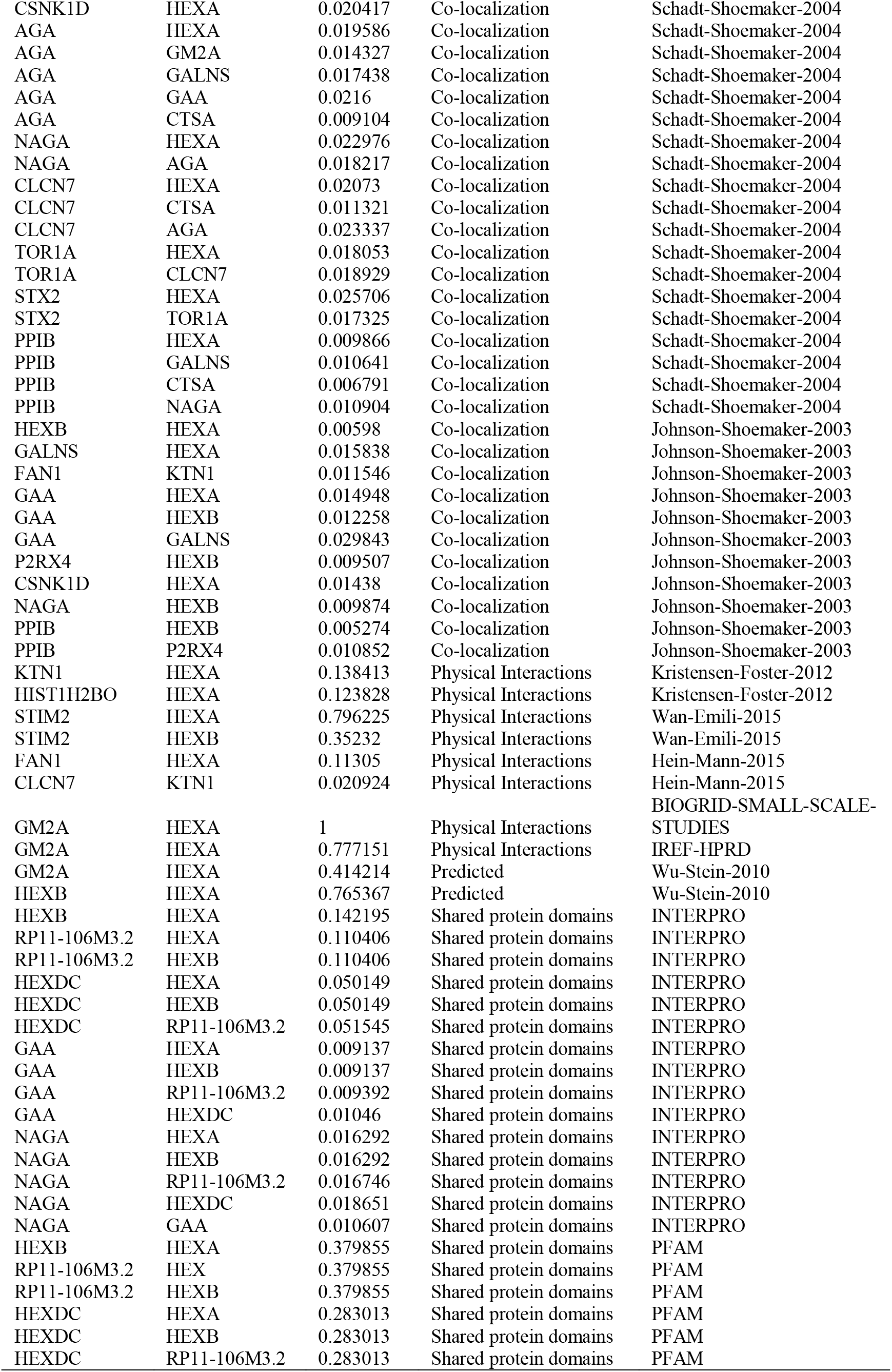
The gene co-expressed, share domain and Interaction with *HEXA* gene network

### SNPs effect on 3’UTR Region (miRNA binding sites) using PolymiRTS Database

**Table (6):**
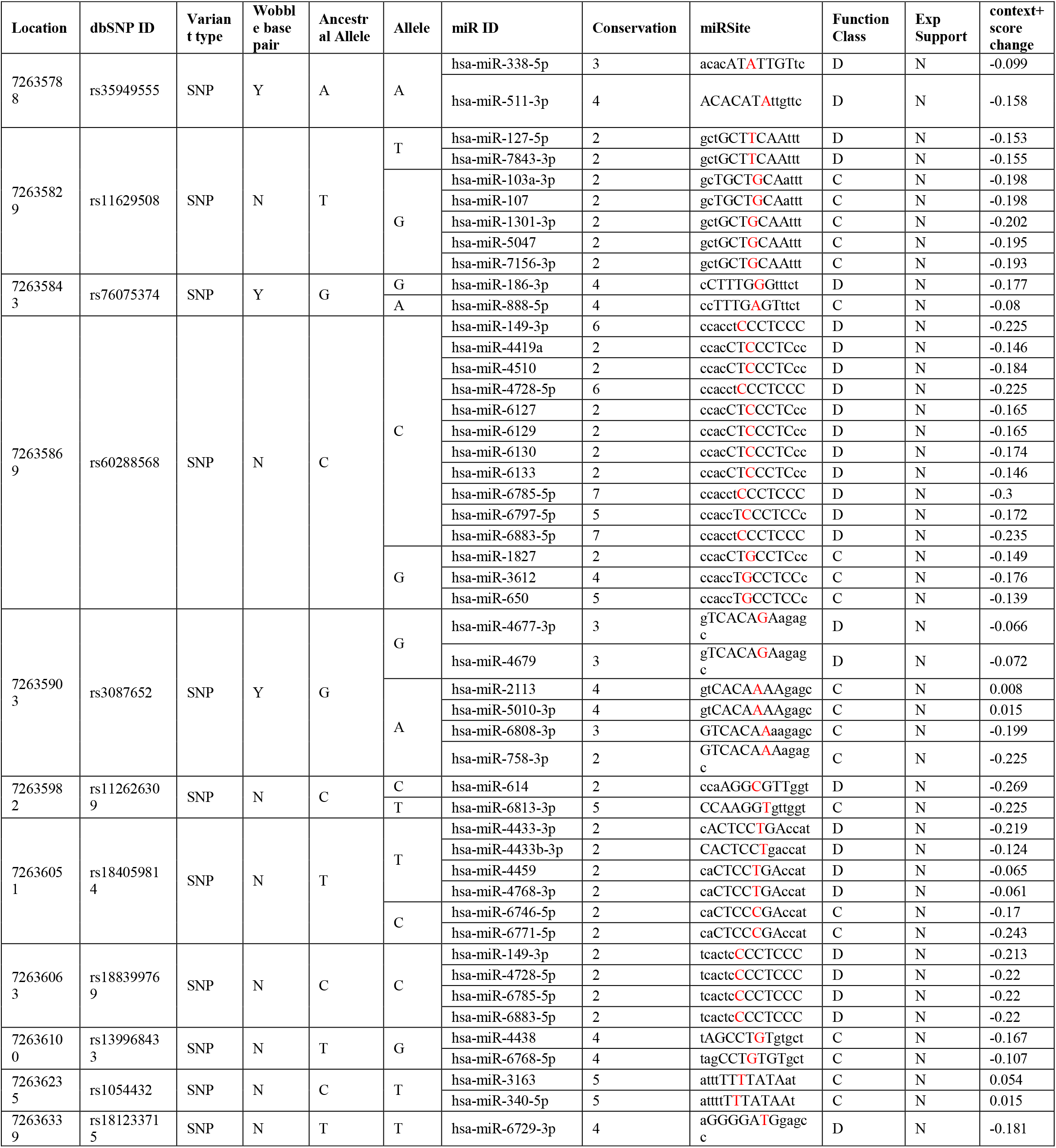
prediction of SNPs at the 3’UTR Region using PolymiRTS

## DISCUSSION

In this study, we revealed 32 novel nsSNP in *HEXA* gene (mentioned in Table (7) out of 59 most damaging missense mutations (in coding region of gene) by using different softwares. We will discuss some of the SNPs that revealed to be most damaging SNPs by this computational analysis study (E462V, R247W, R178H, and R170W).

E462V (1385A>T) mutation was identified in different Indian populations and mentioned to be one of the most commonly reported missense mutations in Indian TSD patient [15, 33, 34]. It was identified as a founder mutation in about 21-40% of TSD patients from Gujarat in India [35–37]. Depending on previous study findings, patients originating from Gujarat state could be screened for E462V mutation [8, 38]. E462V firstly described in Mistri et al. (2012) study as a novel mutation found in six unrelated families from Gujarat indicating a founder effect. They found that it is occurring at the functionally active site of the alpha subunit of hexosaminidase A. In their analysis they found that E462 is a highly conserved region. This mutation was present in the homozygous state in all six patients exhibiting infantile TSD. This is in accordance with previous observations that missense mutations responsible for infantile TSD were generally located in a functional importance region, such as the active site. Also, they recommend screening of E462V SNP in TSD patients from Gujarat [8] and that is in agreement with this study finding in that this SNP is predicted to be a damaging mutation.

R247W (739C>T) accompanied by R249W (745C>T) regarded in Cao, Z., et al.(1997) study as benign mutations that reduce the alpha subunit of protein through affecting its stability in vivo without affecting its processing i.e. phosphorylation, targeting or secretion. Studies also demonstrated that these benign mutations could be readily differentiated from disease-causing mutations using a transient expression system[39].

R247W(739C>T) together with R249W (745C>T) occur in 2% of Ashkenazi Jewish and in 35% in non-Jews and considered to be pseudodeficient mutations, because of that the two common variations of the *HEXA* gene lead to reduced activity against the artificial MUG (4-methylumbelliferyl-2-acetamido-2-deoxy-D-glucopyranoside) substrate but do not impair the ability to degrade the physiological substrate GM2,sothey are associated with no clinical phenotypes [8, 15, 40–44]. These results disagree with this study which record that these SNPs are causing disease and effect on protein structure, function and stability by using different tools to prove that.

R170W (508C>T) mutation identified in previous studies to be associated with an infantile form of TSD [19, 34, 45–47]. It shows complete inactivation of Hex A by the consistent with the infantile TSD phenotype. In areported case, the R170W substitution is associated with the expression of a subunit precursor while no association with its maturation and targeting to the lysosome [46]. R170W was one of the amino acid substitutions which have been reported with a study in the alpha subunit of Hex A. The dysfunctional and destabilizing defects in Hex α- reflects biochemical and phenotypic abnormalities in TSD [47]. In another study, R170W identified as a known pathogenic mutation which disrupts the beta sheet. This mutation has been previously reported in different ethnic groups [Japanese, French-Canadian (Estreeregion, Quebec) and Italian patients]. The side chain of R170 in domain II forms a hydrogen bond with the E141 in domain I. The substitution of R with W with a bulkier side chain has a large effect on the interface of domains I and II. This would destabilize the domain interface and cause degradation of the a-subunit[8].So they agree with this study result in that R170W SNP consider as harmful mutation.

R178H(533G>A) is the most reported mutation responsible for the TSD B1 variant[45, 48–53]. It is mainly found in a higher frequency in Portugal [45, 48–50, 52, 54, 55]. It found in *HEXA* gene and affect the active site residue in the alpha subunit of Hex A enzyme, affecting its structure and subsequently its function by inactivating it leading to Tay-Sachs disease [45, 47, 49, 51, 54–56]. And that is confirming this study outcome to be a damaging SNP. Using I-Mutant software, 12SNPs were predicted to increase the stability of protein while 47 SNPs were predicted to decrease it.

Out of the 154 SNPs in 3’UTR, 11SNPs were found to have an effect on it. 48 functional classes were predicted among 11 SNPs; 28 alleles disrupted a conserved miRNA site and 20 derived alleles creating a new site of miRNA and this might result in deregulation of the gene function. GeneMANIA revealed *HEXA* gene functions and activities such which they are following: aminoglycan catabolic process, glycolipid metabolic process, glycosaminoglycan catabolic process, glycosphingolipid metabolic process, keratan sulfate catabolic process, keratan sulfate metabolic process, lysosomal lumen, membrane lipid metabolic process, sphingolipid metabolic process, sulfur compound catabolic process, vacuolar lumen, and vacuolar part. As bioinformatics softwares play an important role, invivo/invitro analysis remains highly recommended confirming these present study findings. These findings can be used in a helpful way to improve the diagnosis of TaySaches Disease.

## CONCLUSION

This study revealed 32 novel nsSNPout of 59 damaging SNPs in the *HEXA* gene that leads to TaySaches disease, by using different bioinformatics tools. Also, 48 functional classes were predicted in11 SNPs in the 3’UTR, among the 21 SNPs, 28 alleles disrupted a conserved miRNA site and 20 derived alleles created a new site of miRNA. This might result in the de-regulation of the gene function. Hopefully, these results will help in genetic studying and diagnosis of TaySaches disease improvement.

## ACKNOWLEDGMENT

The authors desire to acknowledge the exciting cooperation of Africa City of Technology – Sudan.

## DATA AVAILABILITY

All relevant data used to support the findings of this study are included within the manuscript and supplementary information files.

